# VESTA: Machine Learning-Enabled Estimation of ViscoElastic Ratios from On-Axis Spatio-Temporal ARFI Features

**DOI:** 10.64898/2026.07.06.736692

**Authors:** Simina Mannan Trisha, Md Ashiqur Rahman, Md Walid Hassan, Young Jin Gi, Jangsoon Lee, Md Murad Hossain

## Abstract

Viscoelastic characterization of tissue has significant diagnostic value in oncology, as tumor progression alters both elasticity and viscosity in ways that neither property alone can fully capture. Existing acoustic radiation force (ARF)-based methods such as Viscoelastic Response (VisR) ultrasound estimate relative elasticity and viscosity through per-A-line nonlinear model fitting, which is computationally intensive and requires auxiliary simulations to correct elasticity-dependent bias. This work presents VESTA (Machine Learning-Enabled Estimation of ViscoElastic Ratios from On-Axis Spatio-Temporal ARFI Features), a two-stage data-driven pipeline that predicts elasticity ratio (ER) and viscosity ratio (VR) directly from seven normalized ARFI displacement features at the A-line level, without model fitting or compensation. Stage 1 is an MLP classifier that detects inclusion boundaries from normalized peak displacement and negative peak velocity ratios; Stage 2 is a dilated Conv1D regression model that estimates ER and VR along the full axial sequence using the predicted mask alongside displacement features. The pipeline was trained on 500 simulated inclusion scenarios spanning three geometries, five focal depths, two F-numbers, and a broad range of material contrasts. In silico, mean predicted ER and VR were within 12% of ground truth across all geometries, with performance best when ER and VR were moderate or decoupled. Experimental validation on a chicken breast phantom demonstrated plausible generalization to real tissue heterogeneity. Applied to an in vivo murine 4T1 breast cancer model, the pipeline tracked treatment-related attenuation of mechanical contrast in paclitaxel-treated tumors relative to controls over a 36-day imaging period, supporting its relevance for tumor monitoring.

## 1 Introduction

Cancer progression is associated with localized changes in tissue microstructure, including increased extracellular matrix deposition, elevated collagen density, and enhanced cellular proliferation, which alter the mechanical properties of the tumor microenvironment [1,2]. These changes affect both tissue elasticity and viscosity; elasticity alone is insufficient for comprehensive tumor characterization, as tissue viscosity has been shown to reflect extracellular matrix composition, cell density, and fluid turnover [3, 4]. Imaging frameworks capable of jointly quantifying both properties are therefore essential for tumor diagnosis and treatment monitoring.

Magnetic resonance elastography (MRE) has emerged as a leading technique for quantitative viscoelastic tissue characterization. MRE applies low-frequency mechanical vibrations to tissue and encodes the resulting shear-wave propagation using motion-sensitized MR sequences, reconstructing spatially resolved maps of both the storage modulus (elasticity) and loss modulus (viscosity) [5]. MRE has demonstrated diagnostic value in breast cancer, prostate cancer, and hepatocellular carcinoma, where viscoelastic parameters have been shown to distinguish malignant from benign lesions and to monitor treatment response [1, 6]. However, MRE requires specialized hardware, motion-encoding MR sequences, dedicated reconstruction software, and high-field MRI scanners, substantially restricting its clinical accessibility [7,8]. These constraints motivate the development of alternative viscoelastic imaging methods on conventional diagnostic ultrasound systems.

Ultrasound-based viscoelastic assessment can be achieved by exploiting acoustic radiation force (ARF), which induces tissue displacements monitored either on-axis within the ARF excitation region or off-axis through shear wave propagation [9]. Off-axis methods such as shear-wave elastography (SWE), shear-wave dispersion ultrasound vibrometry (SDUV), and Supersonic Shear Imaging (SSI) estimate elasticity from shear-wave speed and viscosity from frequency-dependent dispersion [10–12], but require accurate lateral wave tracking and are sensitive to noise, attenuation, and boundary conditions in heterogeneous media. On-axis ARF-based methods analyze displacement along the excitation beam axis, improving spatial localization and reducing sensitivity to shear wave reflections [13, 14], though they are sensitive to depth-dependent ARF amplitude variation and often yield only qualitative contrast [15]. Early on-axis approaches include ARFI imaging [13, 14], MSSER [16], harmonic motion imaging (HMI), and ARF-induced creep-recovery (ARFICR) [17], as well as model-based parameter estimation methods that fit displacement data to rheological models but are sensitive to noise and model assumptions.

The Viscoelastic Response (VisR) framework was developed to enable efficient viscoelastic assessment by fitting on-axis displacement from two co-localized ARF excitations to a mass–spring–damper (MSD) model, yielding relative elasticity (RE) and relative viscosity (RV) [18–20]. VisR has been applied to renal transplant evaluation [21], skeletal muscle characterization [22], and breast cancer diagnosis [23]. However, RE and RV estimation requires nonlinear MSD fitting for every A-line, which is time consuming and computationally intensive [9, 20]. Furthermore, RV is confounded by tissue stiffness due to complex system inertia, requiring additional FEM simulations for correction [15, 20], and even after correction RV does not scale proportionally with true viscosity [20]. These limitations motivate alternative approaches that are scalable and less model-dependent.

Machine learning has been increasingly applied to viscoelastic property estimation from ARF-induced displacement data [24, 25]. Richardson et al. [26] proposed Quantitative VisR (QVisR), a fully-connected neural network trained on FEM and Field II simulations that estimates absolute elastic and viscous moduli from VisR displacement profiles, with earlier work demonstrating feasibility in homogeneous materials [27]. However, these approaches operate on full displacement time-series in homogeneous settings and do not address spatially resolved inclusion detection or A-line-level prediction across heterogeneous geometries with varying focal depths.

This work proposes a data-driven framework for estimating elasticity ratio (ER) and viscosity ratio (VR) at the A-line level from ARFI-derived displacement features. Finite Element Method (FEM) and Field II acoustic simulations generate labeled training data with known mechanical properties. A two-stage pipeline — comprising an MLP classifier for inclusion detection and a dilated Conv1D regression model for ER and VR prediction — is evaluated on unseen simulated materials and an experimental tissue-mimicking phantom. The objectives are: (1) to assess whether normalized displacement features capture viscoelastic contrast across heterogeneous geometries; (2) to evaluate ER and VR prediction across varying material contrasts; and (3) to assess framework robustness using simulation-based datasets.

## 2 Methodology

### 2.1 In Silico Model

Viscoelastic inclusion phantoms were simulated using a FEM model adapted from [20], solved via LS-DYNA3D (Livermore Software Technology Corporation, CA, USA) with an explicit time-domain method.

A 3-D rectangular mesh of 0.25 mm^3^ cubic elements was assembled using LS-PREPOST. Two mesh configurations were used depending on inclusion size: the smaller mesh spanned 3–43 mm axially and ± 8 mm laterally; the larger mesh extended laterally from −8 to +11 mm. Elevational dimensions of ±6 mm (mesh) were identical for both configurations.

Tissue was modeled as an incompressible isotropic viscoelastic material using the MAT_KELVIN _MAXWELL_VISCOELASTIC (MKMV). The MKMV model consists of a spring (*µ*_*o*_) in series with a Voigt unit comprised of a spring (*µ*_*α*_) and a dashpot (*η*) in parallel. To approximate Voigt material behavior, *µ*_*o*_ was set to 200 times the value of *µ*_*α*_, consistent with prior VisR simulations [20]. Material density was 1000 kg m^*−*3^ and Poisson’s ratio was 0.499. A five-element-thick perfect matching layer (PML) suppressed spurious boundary reflections.

### 2.2 Inclusion Geometry and Dataset Design

Three geometries were simulated: spherical, cuboidal, and ellipsoidal. Spherical and cuboidal inclusions had diameters or side lengths of 3 mm and 6 mm. Ellipsoidal inclusions were oriented at 90 degrees in four size variants: semi-minor axis 2 mm with semi-major axes of 3 mm and 5 mm, and semi-minor axis 4 mm with semi-major axes of 5.5 mm and 7.5 mm. The background A-line was fixed at 6.0 mm lateral for the smaller mesh and 9.0 mm for the larger mesh.

The dataset comprised 500 inclusion scenarios across six focal depths (15, 19, 20, 23, 27, 30 mm), with around 50 to 100 inclusions per depth balanced as 34 spherical, 33 cuboidal, and 33 ellipsoidal. Four lateral A-line indices were sampled per inclusion (3 inclusion lines and 1 background line), yielding 2000 total simulations.

### 2.3 Material Parameter Sampling

Background shear elastic moduli (*G*_*b*_) and viscosities (*µ*_*b*_) were sampled log-uniformly from 1–10 kPa and 0.1–5 Pa·s. Inclusion properties were defined via *r*_*G*_ = *G*_*i*_*/G*_*b*_ and *r*_*µ*_ = *µ*_*i*_*/µ*_*b*_, both constrained to [1.1, 16] and [1.1, 10] respectively for ER and VR, with absolute bounds *G*_*i*_ ∈ [1, 50] kPa and *µ*_*i*_ ∈ [0.1, 10] Pa·s enforced by rejection sampling.

Ratios were sampled log-uniformly across four contrast tiers: weak (1.1–2.0, 25%), medium (2.0–4.0, 35%), strong (4.0–7.0, 25%), and ultra (7.0–16.0, 15%). To avoid spurious coupling between elasticity and viscosity, three decoupling groups were defined: Group A (*n* = 300, both *r*_*G*_ and *r*_*µ*_ vary freely), Group B (*n* = 100, elasticity-dominant: *r*_*µ*_ *∈* [1.1, 1.3]), and Group C (*n* = 100, viscosity-dominant: *r*_*G*_ *∈* [1.1, 1.3]).

### 2.4 ARF Field and Ultrasonic Tracking Simulation

The ARF excitation field was computed at each FEM node using Field II [33] following the method of Palmeri et al. [32], with L11-5 transducer parameters. Intensity values were scaled to 5000 W/cm^2^ and the volumetric ARF magnitude was computed as [3]:

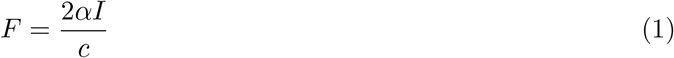

where *α* = 0.5 dB cm^*−*1^ MHz^*−*1^, *c* = 1540 m/s, and *I* is the temporal-average beam intensity. Point forces of duration 70 *µ*s were applied to the mesh. Simulation duration was 3 ms with data saved every 0.067 ms (PRF = 15 kHz) [14].

Ultrasonic tracking of ARF-induced displacements was simulated following the approach of Palmeri et al. [34]. A 3-D Field II scatterer phantom with fully developed speckle (15 scatterers/resolution cell) was defined to span the FEM mesh volume. Two push F-numbers, 2.0 and 2.5, were simulated, with corresponding tracking F-numbers of 1.5 and 2.0, respectively. FEM-derived displacements were used to linearly interpolate scatterer positions at each time step in the ARFI ensemble using MATLAB (MathWorks, Inc., Natick, MA, USA). After generating scatterer position matrices for each time step, the corresponding RF lines were simulated using Field II with the L11-5 transducer tracking parameters described above. White Gaussian noise was added to simulate a system echo SNR of 35 dB. Motion tracking was performed using one-dimensional normalized cross-correlation (NCC) [35], yielding axial displacement time-profiles at each simulated A-line location.

### 2.5 Displacement Feature Extraction and Normalization

Displacement waveforms were processed offline in MATLAB (MathWorks, Natick, MA) through sequential bandpass filtering, axial moving mean smoothing, and linear detrending to suppress noise and bulk motion. Seven displacement-derived features were extracted at every axial depth: peak displacement (PD), positive peak velocity (PV^+^), negative peak velocity (PV^*−*^), time to peak displacement (TTP), time to 80% of peak displacement (TTP_80_), time to 60% of peak displacement (TTP_60_), and time to negative peak velocity (TPV). These features collectively characterize displacement magnitude, recovery dynamics, and tissue velocity response during ARF loading and relaxation.

In on-axis ARF methods, the radiation force field is spatially non-uniform due to beam focusing and depth-dependent attenuation [14,15], and impedance mismatches at inclusion boundaries further alter the effective force distribution inside the inclusion relative to the background [9]. Each feature was therefore normalized by its background counterpart at the same axial depth:

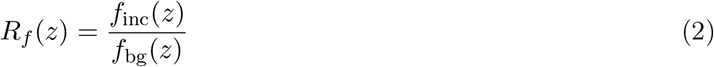

where *f*_inc_(*z*) and *f*_bg_(*z*) denote a given feature at axial depth *z* for the inclusion and background A-lines, respectively. This yields seven normalized ratio features: 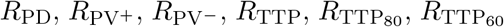, and *R*_TPV_.

### 2.6 Summary of Parameter Ranges

The test materials were selected with mechanical properties distinct from those used during training; however, acoustic properties (speed of sound = 1540 m/s, attenuation = 0.5 dB cm^*−*1^ MHz^*−*1^) and ARFI push and tracking parameters (F-number = 2.5, focal depth = 20 mm) were identical between training and test groups.

### 2.7 Machine Learning Pipeline

A two-stage machine learning pipeline was developed to estimate ER and VR at the A-line level from the normalized feature profiles described above. Stage 1 is a binary classifier that predicts an inclusion mask from displacement features, and Stage 2 is a regression model that predicts ER and VR using the predicted mask alongside the displacement features. Training of these two stages involved four sequential steps: Step 1 trained Stage 1 via 5-fold cross-validation to generate out-of-fold mask predictions; Step 2 used those mask predictions alongside the displacement features to train Stage 2; Step 3 retrained a final Stage 1 model on the full training set; and Step 4 evaluated the complete pipeline on a held-out validation set. The full dataset was partitioned into a training set

#### 2.7.1 Step 1: Stage 1 training with 5-Fold cross validataion

Stage 1 was formulated as binary classification, with ground-truth labels of 1 inside the inclusion and 0 in the background. *R*_PD_ and 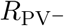 were used as input features, standardized using the mean and standard deviation from the training subset of each fold.

The classifier was a fully connected neural network with three hidden layers of 256, 128, and 64 neurons, each with batch normalization and ReLU activation. Dropout (*p* = 0.2) was applied after the first two hidden layers and L2 regularization was applied throughout. The output layer used sigmoid activation, thresholded at 0.5 for binary prediction. The network was trained with binary cross-entropy loss using the Adam optimizer (learning rate 3 × 10^*−*4^), a batch size of 512, and up to 250 epochs.

A five-fold cross-validation strategy was used with all splits performed at the material-case level. In each fold, 20% of materials formed the out-of-fold validation set, among the 80%, 12% were reserved as an internal validation set for early stopping, and the remaining 68% were used for epoch-by-epoch training. This process was repeated across all five folds to generate out-of-fold mask predictions for the full training set, which were subsequently used as Stage 2 inputs.

#### 2.7.2 Step 2: train Stage 2 to predict ER and VR

Stage 2 received the full axial feature sequence and predicted ER and VR at every axial depth. All seven normalized features (*R*_PD_, 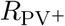, 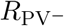, *R*_TTP_, *R*_TTP80_, *R*_TTP60_, *R*_TPV_) were used alongside the Stage 1 predicted mask 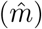, focal depth (FD), normalized axial depth (*z*_norm_ = (*z−* FD)*/*FD), and F-number, forming an 11-feature input vector at each axial location. Features were standardized using statistics from the training sequences. Axial samples classified as background by Stage 1 were assigned ER = 1 and VR = 1 prior to training so that the model learned to predict reference values in background regions.

The Stage 2 model was a dilated Conv1D network operating on sequences of size *N*_*z*_ × 11. Three Conv1D layers extracted spatial patterns along the axial direction: 32 filters with kernel size 3 and dilation rate 1; 64 filters with kernel size 5 and dilation rate 3; and 128 filters with kernel size 5 and dilation rate 6. All convolutional layers used L2 regularization, batch normalization, and softplus activation. Three subsequent time-distributed dense layers of 256, 128, and 64 neurons with L2 regularization, batch normalization, softplus, dropout *p* = 0.2 after the first two further refined the representation. The output layer produced [ER, VR] at every axial sample.

Training used a custom weighted mean percentage error (MPE) loss averaged over ER and VR:

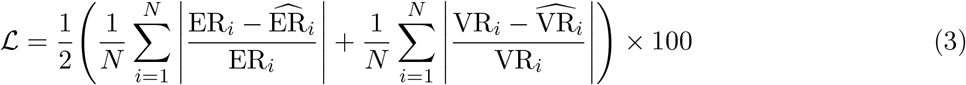

The Adam optimizer (learning rate 3 ×10^*−*4^), batch size of 64, and up to 500 epochs with early stopping were used. 15% of the materials were reserved for validation.

#### 2.7.3 Step 3: Stage 1 Retraining

Following cross-validation, a final Stage 1 model was retrained on all available training data using the same architecture, input features, optimizer, and early-stopping procedure described in Stage 1. The final model weights and feature scaler were saved for use in Stage 4 inference.

#### 2.7.4 Step 4: Full Testing Pipeline and Evaluation

The trained Stage 1 and Stage 2 models were applied sequentially to unseen test cases. Stage 1 predicted the inclusion mask for each A-line, which was then passed as input to Stage 2. Stage 2 predicted ER and VR at every axial sample, with background locations assigned ER = 1 and VR = 1 in the final output.

Stage 1 performance was assessed using accuracy, precision, recall, and F1-score. Stage 2 regression performance was evaluated using mean percentage error (MPE) computed separately for ER and VR over inclusion-region axial samples:

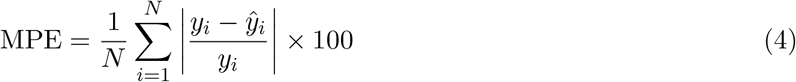

where *y*_*i*_ and *ŷ*_*i*_ are the ground-truth and predicted values, respectively.

### 2.8 Chicken Breast Phantom Preparation

Gelatin-based tissue-mimicking background material was prepared following established phantom recipes, in which gelatin concentration sets the target stiffness and graphite concentration sets the acoustic attenuation [20]. For a background stiffness of 10 kPa and attenuation of 0.5 dB cm^*−*1^ MHz^*−*1^, gelatin was presoaked in water and n-propanol, heated to 65 °C, mixed with graphite, and cooled to 28 °C under stirring before solidifying at room temperature and refrigeration.

A trimmed chicken breast specimen was embedded as an inclusion at a depth of 19 mm within the solidified 10 kPa gelatin background. The phantom was imaged within 48 hours of fabrication at a focal depth of 19 mm using the acquisition protocol described in Section 2.4.

### 2.9 In Vivo Murine Tumor Model

A syngeneic 4T1 triple-negative breast cancer mouse model was used for in vivo validation of the proposed framework. A suspension of 1 × 10^5^ 4T1 cells in 50 *µ*L phosphate-buffered saline (PBS) was injected subcutaneously into the right mammary fat pad of female BALB/c mice. Animals were assigned to a control group or a paclitaxel-treated group. ARFI imaging was performed from Day 2 to Day 33 post implantation in the same animals to capture the temporal evolution of tumor elasticity ratio (ER) and viscosity ratio (VR).

For imaging, Verasonics Vantage NXT ultrasound system with L11-5 transducer was used and mice were placed in a supine position under isoflurane anesthesia (3–5% induction, 1.5–2% maintenance) delivered via nose cone. The tumor-bearing region was depilated and ultrasound coupling gel was applied. ARFI excitations were delivered at a focal depth of 18 mm. Displacement features were extracted, normalized, and processed using the same pipeline described in Section 2.5, and the trained Stage 1 and Stage 2 models were applied to generate predicted ER and VR maps at each imaging timepoint. All animal procedures were conducted in accordance with IACUC protocols.

## 3 Results

### 3.1 Qualitative Assessment of Predicted ER and VR Images

Fig. 1, show ground-truth and predicted ER and VR images for spherical, ellipsoidal, and cuboidal inclusions, respectively, all simulated with the same material contrast with inclusion: *G*_*i*_ = 20 kPa, *µ*_*i*_ = 4 Pa·s; background: *G*_*b*_ = 5 kPa, *µ*_*b*_ = 1 Pa· s; ER = 4, VR = 4, allowing direct comparison of prediction performance across inclusion geometry independent of material contrast. Fig. 2 shows another ellipsoidal inclusion with a different, decoupled material contrast of inclusion: *G*_*i*_ = 8.7 kPa, *µ*_*i*_ = 1.2 Pa·s; background: *G*_*b*_ = 1.4 kPa, *µ*_*b*_ = 0.8 Pa·s; ER = 6.2, VR = 1.5, enabling assessment of the model under a higher-ER, lower-VR contrast regime. For all four cases, the predicted parametric images qualitatively delineate the inclusion boundary against the homogeneous background, with predicted ER and VR values elevated within the inclusion region and returning to the reference background value of unity outside the inclusion, closely matching the ground-truth distribution.

**Figure 1.**
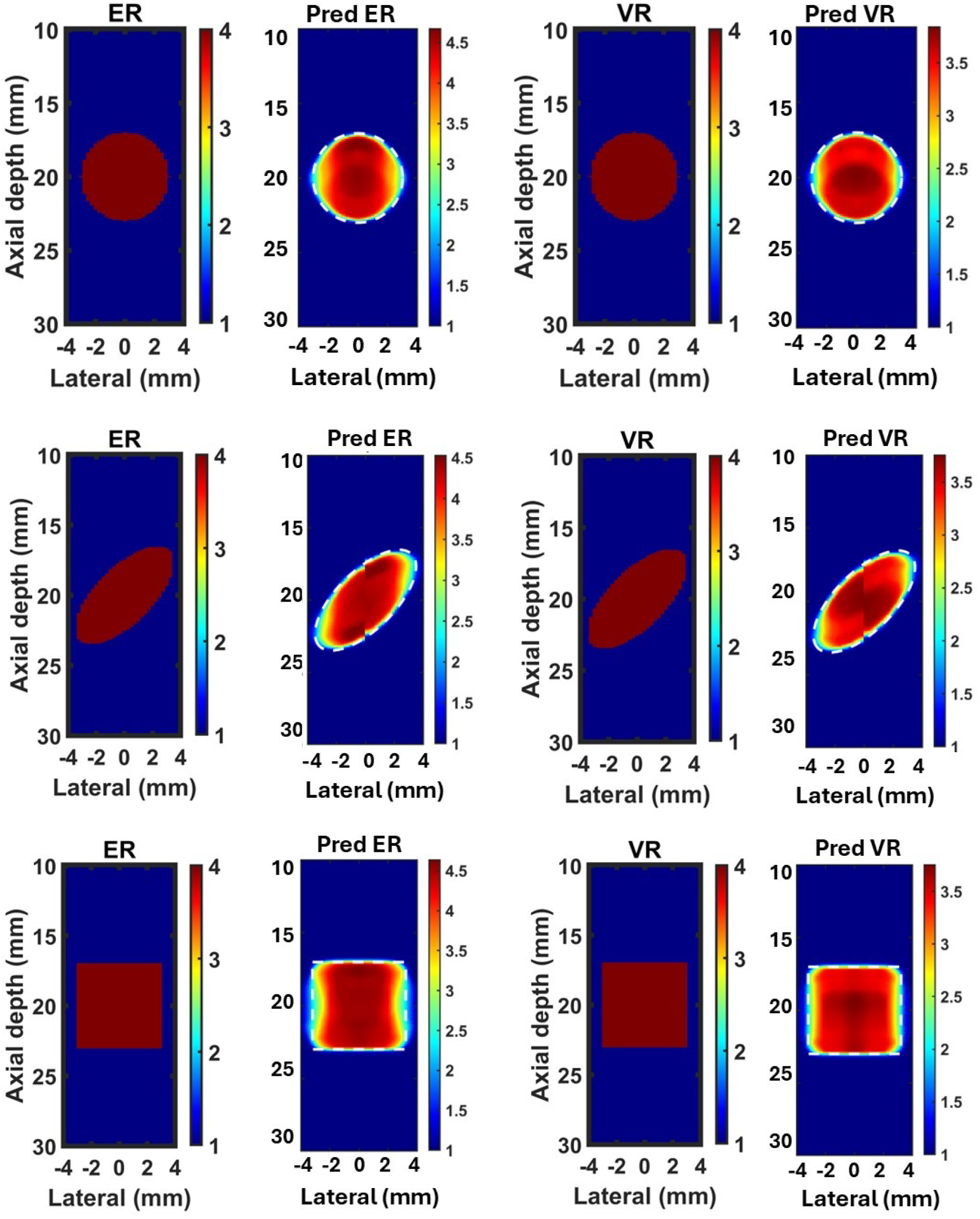
Ground-truth and predicted ER and VR images for three inclusion geometries of identical material contrast (Inclusion:*G*_*i*_ = 20 kPa, *µ*_*i*_ = 4 Pa· s; Background: *G*_*b*_ = 5 kPa, *µ*_*b*_ = 1 Pa· s; ER = 4, VR = 4). a) ER ground truth b) ER predicted, c) VR ground truth, d)VR predicted.

**Figure 2.**
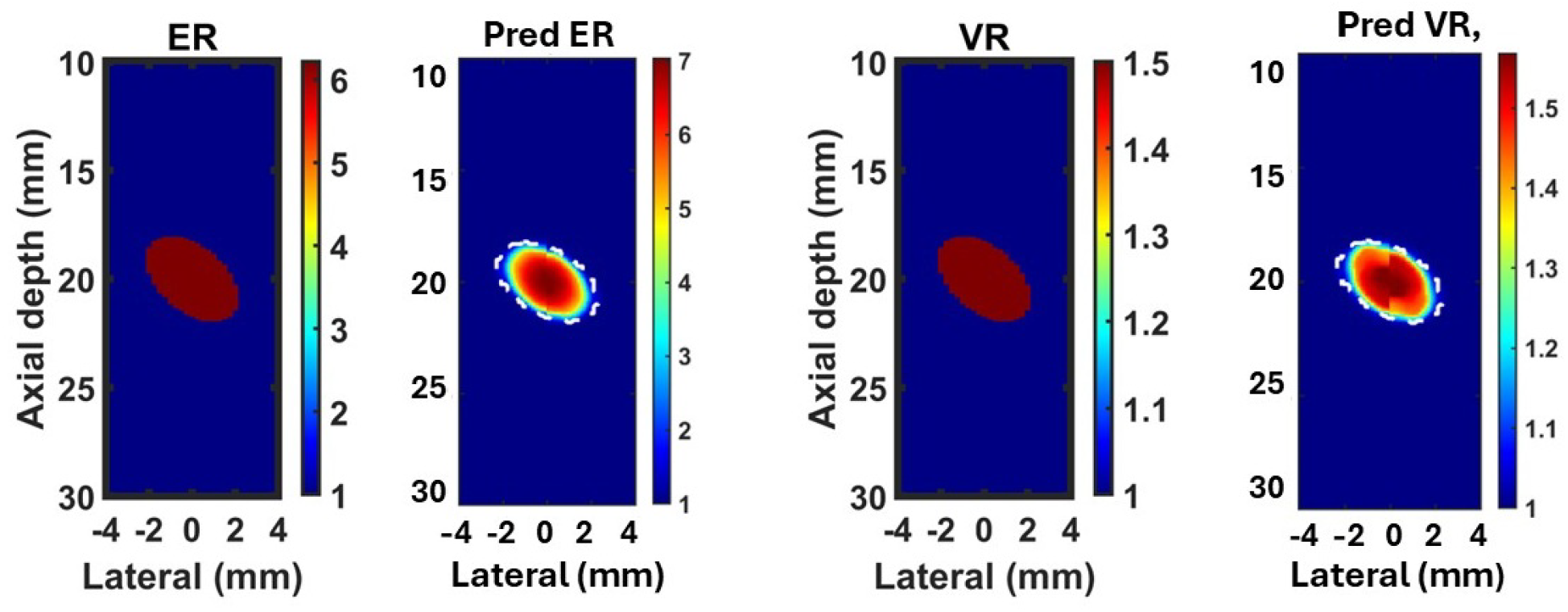
Ground-truth and predicted ER and VR images, in order, for an ellipsoidal inclusion with decoupled material contrast (inclusion: *G*_*i*_ = 8.7 kPa, *µ*_*i*_ = 1.2 Pa· s; background: *G*_*b*_ = 1.4 kPa, *µ*_*b*_ = 0.8 Pa·s; ER = 6.2, VR = 1.5: (a) ER ground truth, (b) ER predicted, (c) VR ground truth, (d) VR predicted.

For the spherical inclusion with high material contrast (ER = 4, VR = 4), the predicted ER and VR images show a well-localized circular region of elevated contrast that closely follows the true inclusion boundary, with mean predicted values of 4.36 for ER and 3.62 for VR, both within 10% of the ground-truth values.

For the ellipsoidal inclusion of the same material contrast, the predicted ER and VR images similarly follow the elongated, obliquely oriented inclusion boundary, with mean predicted values of 4.28 for ER and 3.56 for VR, corresponding to relative errors of 7.0% and 11.0% from the ground-truth ER = 4 and VR = 4. Relative to the spherical inclusion, VR was more underestimated for the ellipsoidal geometry, consistent with the elongated, tilted shape introducing greater boundary-averaging effects.

For the cuboidal inclusion, the predicted ER and VR images show elevated contrast well-contained within the true inclusion boundary, with mean predicted values of 4.46 for ER and 3.58 for VR. The predicted contrast region approximates the square cross-section of the true inclusion, though with rounded corners relative to the sharp-edged ground-truth boundary, consistent with the spatial smoothing introduced by the axial and lateral extent of the ARF excitation and displacement tracking.

For the ellipsoidal inclusion with decoupled material contrast (*G*_*i*_ = 8.7 kPa, *µ*_*i*_ = 1.2 Pa· s; *G*_*b*_ = 1.4 kPa, *µ*_*b*_ = 0.8 Pa s; ER = 6.2, VR = 1.5), the predicted images show mean values of 5.80 for ER and 1.47 for VR, corresponding to relative errors of 6.7% for ER and 1.7% for VR. Both errors are substantially lower than the equal-contrast ellipsoidal case, indicating that the model performs more accurately when ER and VR are decoupled rather than simultaneously elevated, with VR estimation remaining particularly robust when viscosity contrast is close to background.

Across all four cases, predicted ER and VR values were within 12% of ground truth, and background regions outside the inclusion boundary returned to the expected reference value of unity in all cases, indicating that the two-stage pipeline reliably suppresses false-positive contrast outside the true inclusion extent, consistent with the quantitative trends examined further in the following section.

### 3.2 Quantitative Error Analysis

Fig. 3 shows the percent error distribution between predicted and ground-truth ER and VR across six representative material and geometry combinations, spanning elasticity ratios (ER) from 2.5 to 6.2 and viscosity ratios (VR) from 1.5 to 4. Median percent error across all cases ranged from approximately 6.5% to 12.5% for ER and 2.5% to 10.5% for VR.

**Figure 3.**
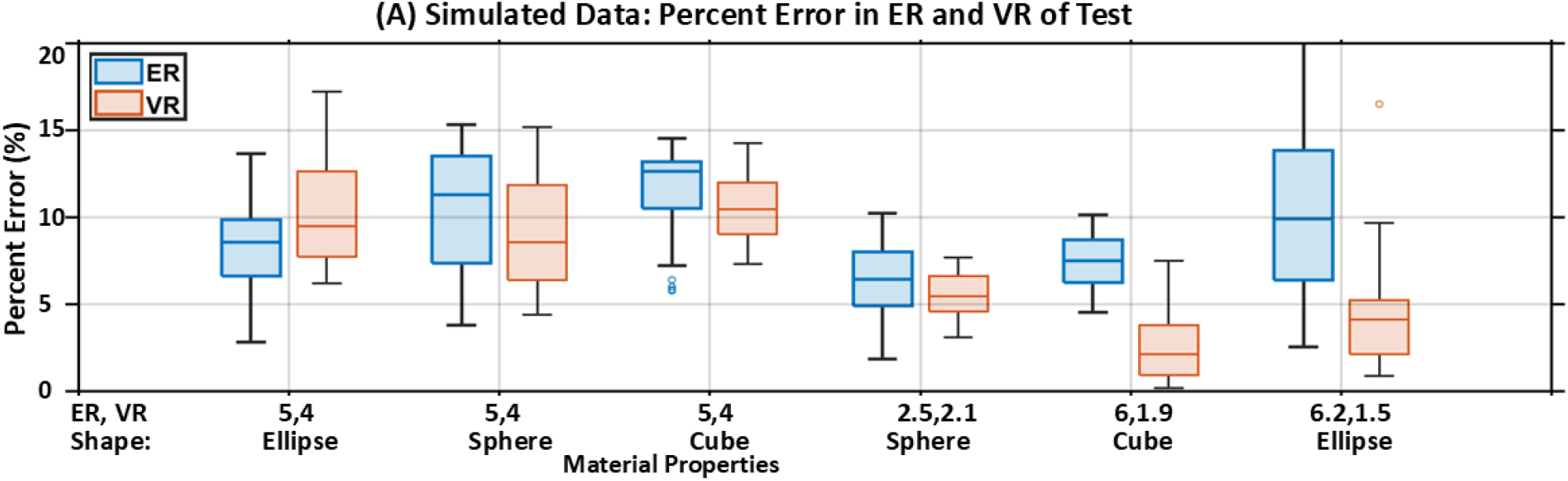
Percent error distributions between predicted and ground-truth ER (blue) and VR (orange) across six representative test cases spanning three inclusion geometries and a range of material contrasts. Ground-truth elasticity ratio (ER) and viscosity ratio (VR) for each case are indicated below the corresponding box pair.

The lowest errors were observed for the cube (*G*_*i*_ = 15 kPa, *µ*_*i*_ = 3.5 Pa· s; *G*_*b*_ = 2.5 kPa, *µ*_*b*_ = 1.8 Pa·s; length = 6 mm; ER = 6, VR = 1.9) and the sphere (*G*_*i*_ = 14 kPa, *µ*_*i*_ = 5.5 Pa· s; *G*_*b*_ = 5.5 kPa, *µ*_*b*_ = 2.6 Pa·s; diameter = 6 mm; ER = 2.5, VR = 2.1), where median VR error fell below 6% and median ER error remained below 8%.

The highest errors were observed for the ellipsoidal inclusion (*G*_*i*_ = 20 kPa, *µ*_*i*_ = 4 Pa· s; *G*_*b*_ = 5 kPa, *µ*_*b*_ = 1 Pa· s; major axis = 9 mm, minor axis = 4 mm; ER = 4, VR = 4) and the cube of the same material (ER = 4, VR = 4, length = 6 mm), where median ER error exceeded 11% for both. Since these two cases share identical material properties but differ in geometry, the elevated error reflects the ER = VR = 4 material combination itself rather than shape.

For the smaller ellipsoidal inclusion (*G*_*i*_ = 8.7 kPa, *µ*_*i*_ = 1.2 Pa· s; *G*_*b*_ = 1.4 kPa, *µ*_*b*_ = 0.8 Pa· s; major axis = 5 mm, minor axis = 3 mm; ER = 6.2, VR = 1.5), ER error exhibited the widest interquartile range among all six cases, while VR error remained comparatively low and tightly distributed.

Across all six cases, VR error was lower than or comparable to ER error in four of six cases, while ER and VR errors were more closely matched for the two high-contrast, equal-ratio cases (ER = VR = 4). These results indicate that the prediction accuracy depends jointly on inclusion geometry and the relative magnitude of ER versus VR, with the model performing most consistently when the two ratios are moderate or clearly decoupled.

### 3.3 Experimental Validation

#### 3.3.1 Chicken Breast Phantom

Fig. 4 shows the B-mode image and predicted ER and VR maps for a chicken breast inclusion embedded in a gelatin background. The predicted maps show elevated ER and VR within the inclusion boundary relative to the surrounding background, with mean± standard deviation of 8.12± 1.83 for ER and 6.70 ±1.26 for VR, indicating that the chicken breast was detected as both stiffer and more viscous than the gelatin background.

**Figure 4.**
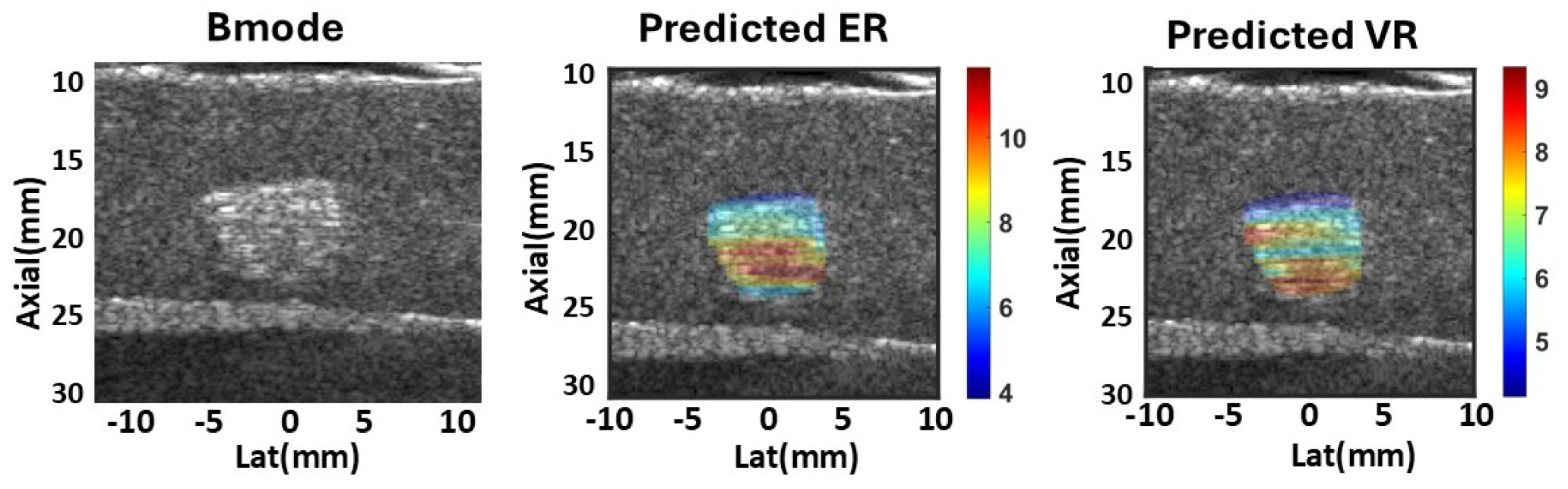
Experimental validation on a chicken breast inclusion embedded in a gelatin background. From left to right: B-mode image, predicted ER image (8.12 ±1.83), and predicted VR image (6.70 ±1.26). White dashed contour indicates the inclusion boundary derived from the B-mode image.

### 3.3.2 Murine Tumor Model

Fig. 5 and Fig. 6 show predicted ER and VR maps, respectively, for representative control and paclitaxel-treated murine tumors at Day 2 and Day 33. In the control tumor, mean ER increased from 2.48 at Day 2 to 7.99 at Day 33, and mean VR increased from 1.82 to 8.45 over the same interval, alongside marked tumor growth visible in the underlying B-mode image. The paclitaxel-treated tumor started from a higher baseline (mean ER = 5.51, mean VR = 3.05 at Day 2) but reached a lower endpoint than control at Day 33 (mean ER = 7.04, mean VR = 5.68), such that the net increase in both ER and VR from Day 2 to Day 33 was substantially smaller in the treated tumor than in control.

**Figure 5.**
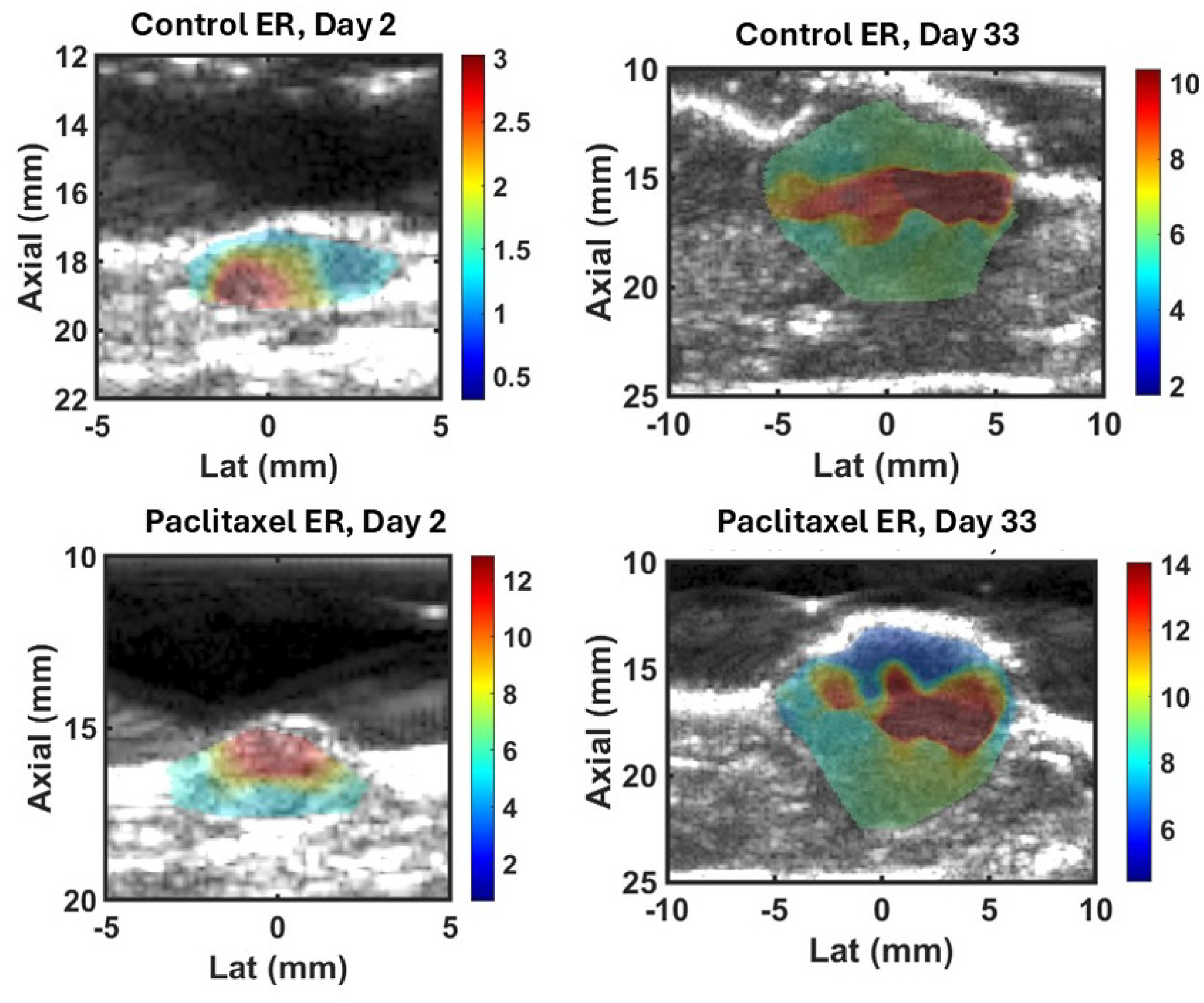
Predicted ER maps for control (top row) and paclitaxel-treated (bottom row) murine tumors at Day 2 (left column) and Day 33 (right column).

**Figure 6.**
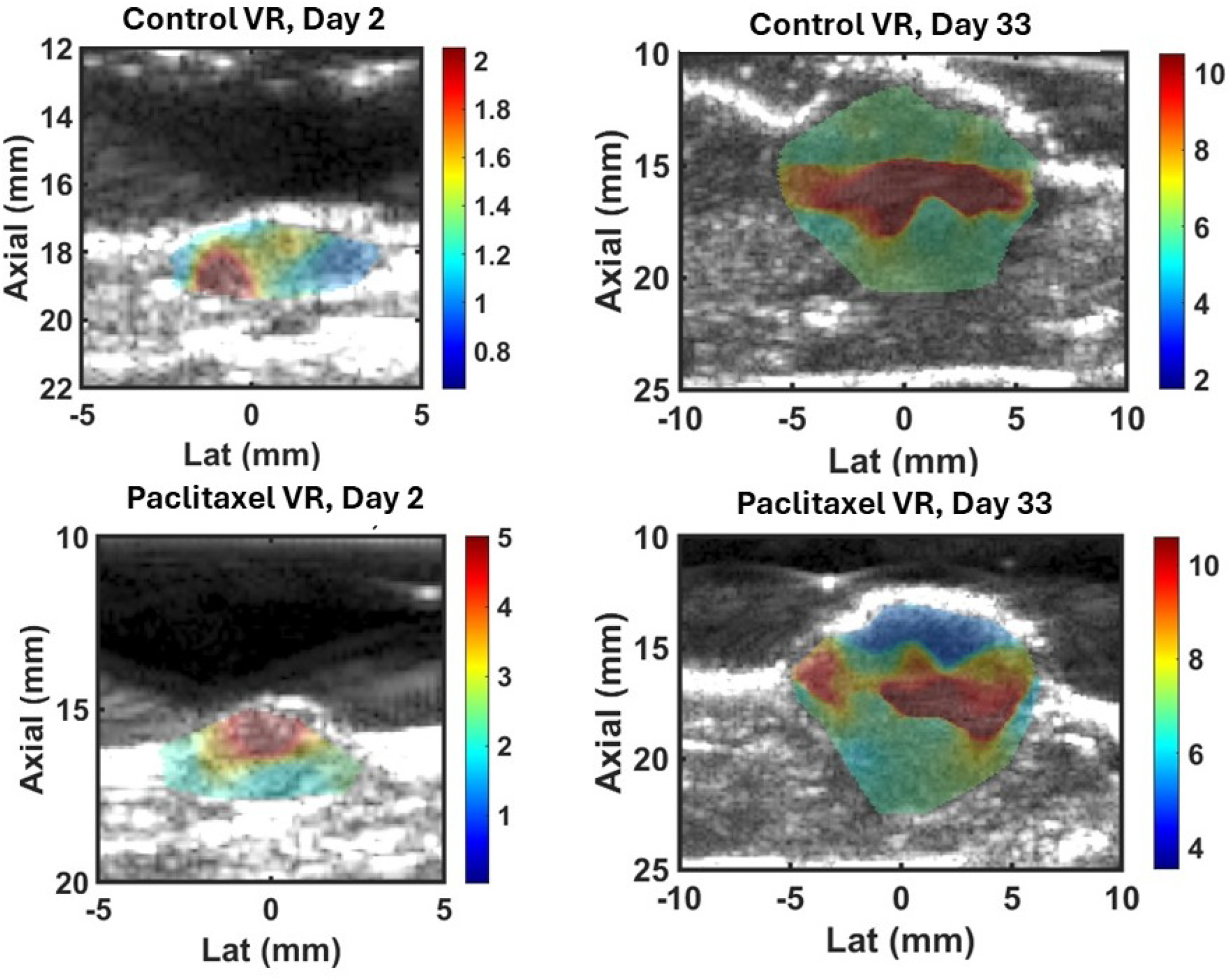
Predicted VR maps for control (top row) and paclitaxel-treated (bottom row) murine tumors at Day 2 (left column) and Day 33 (right column).

Fig. 7 and Fig. 8 show ER and VR distributions, respectively, across all animals at each imaging timepoints from Day 4 to Day 36. Both control and paclitaxel groups exhibit a broadly increasing trend in median ER and VR from Day 4 through a peak near Day 30–32, followed by a decline by Day 36. At Day 4, median ER and VR were similar between groups (ER: 5.1 vs. 5.2; VR: 4.1 vs. 3.1), with overlapping interquartile ranges. From Day 30 onward, however, control values consistently exceeded paclitaxel values in both median and upper quartile, most notably at Day 32 (ER: 9.0 vs. 6.8; VR: 8.1 vs. 5.5), where the paclitaxel group’s interquartile range also widened substantially, indicating greater inter-animal variability in treatment response at this timepoint. This population-level divergence between groups at later timepoints, despite comparable starting values at Day 4, is consistent with the representative-animal trend in Fig. 5 and Fig. 6, in which paclitaxel treatment was associated with an attenuated rise in ER and VR relative to control over the course of tumor growth.

**Figure 7.**
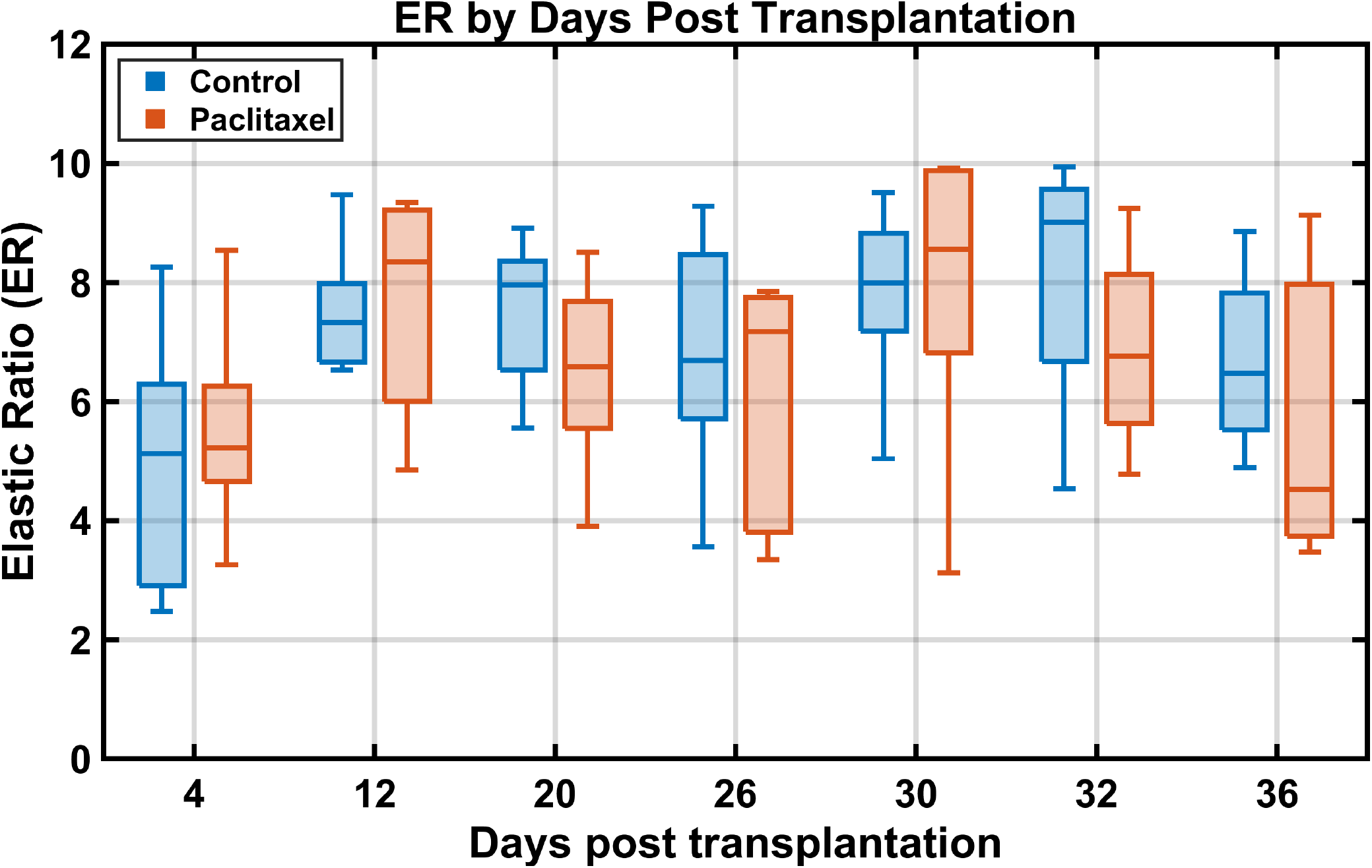
Elastic ratio (ER) distributions for control and paclitaxel-treated murine tumors across imaging timepoints from Day 4 to Day 36 post-transplantation.

**Figure 8.**
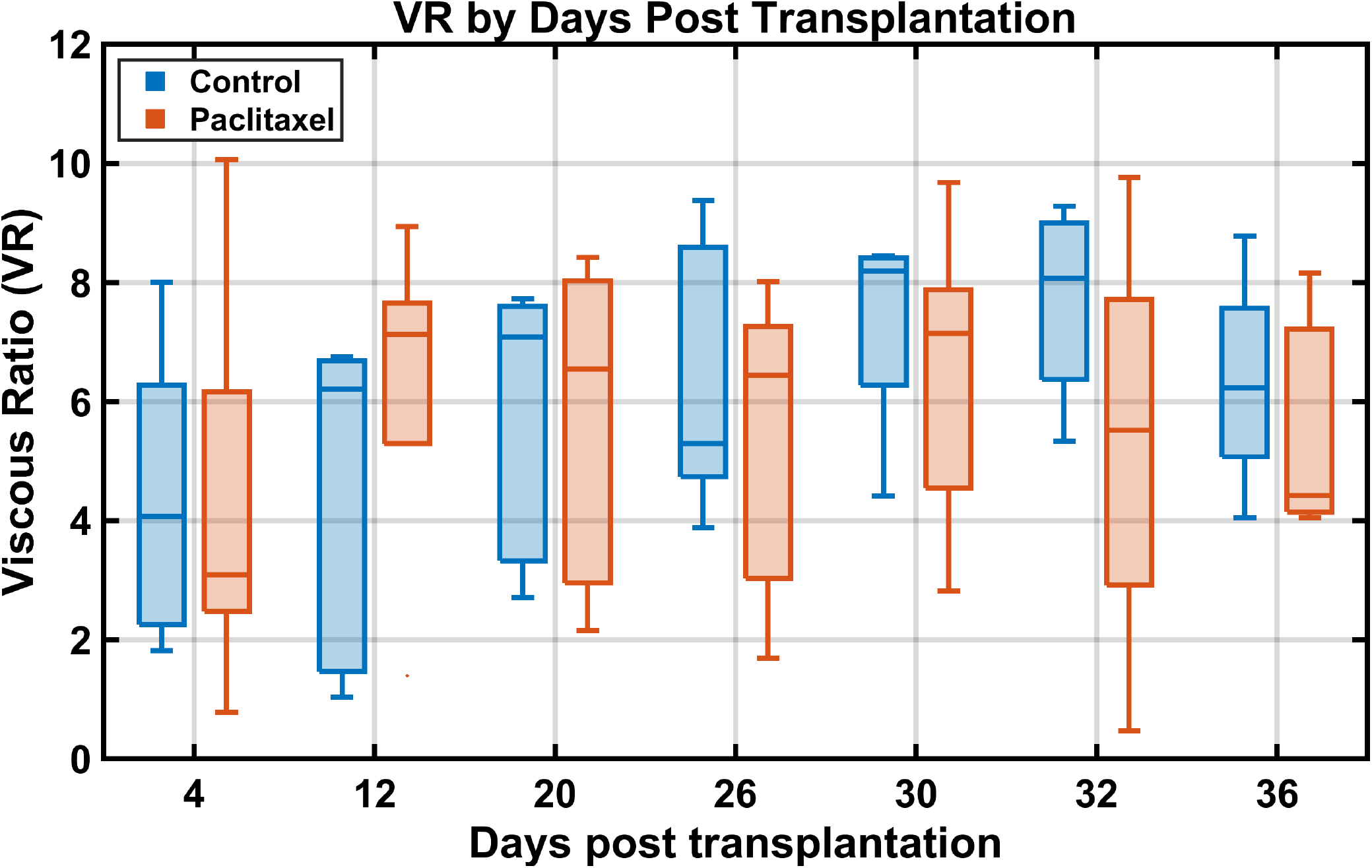
Viscous ratio (VR) distributions for control and paclitaxel-treated murine tumors across imaging timepoints from Day 4 to Day 36 post-transplantation.

## 4 Discussion

The proposed framework is a data-driven method for assessing the viscoelastic properties of tissue by mapping normalized ARFI-induced displacement features directly to elasticity ratio (ER) and viscosity ratio (VR) at the A-line level. Because ER and VR are predicted from a trained model rather than obtained through per-A-line nonlinear MSD fitting, the method avoids the principal time consuming issue and computational bottleneck of VisR-based estimation and does not require the auxiliary FEM simulations that VisR uses to correct the elasticity-dependent bias in RV [20]. Moreover, because the Stage 1 classifier explicitly identifies the inclusion boundary before Stage 2 regression is applied, ER and VR are predicted with spatial specificity rather than as single point measurements, which is expected to better preserve small mechanical features than approaches that rely on spatial averaging over regions of interest.

This work used both FEM simulations and experimental tissue to demonstrate that the proposed pipeline recovers material elasticity and viscosity contrast across heterogeneous inclusion geometries. In Fig. 1, in silico results show that predicted ER and VR closely tracked the true inclusion boundary for all three geometries at identical material contrast (ER = VR = 4). For the spherical and cuboidal inclusions, mean predicted ER and VR were within 12% of ground truth (sphere: ER = 4.36, VR = 3.62; cube: ER = 4.46, VR = 3.58), with errors comparable in magnitude between the two properties. For the ellipsoidal inclusion of the same material, mean predicted values were ER = 4.28 and VR = 3.56, corresponding to errors of 7.0% and 11.0%, respectively, indicating that VR is more sensitive to the elongated, tilted geometry than ER, likely due to greater boundary-averaging effects along the minor axis. Fig. 2 further showed that when ER and VR were decoupled (ER = 6.2, VR = 1.5), both errors dropped substantially relative to the equal-contrast cases (ER = 5.80, error = 6.7%; VR = 1.47, error = 1.7%), suggesting that the network performs more accurately when the two ratios diverge, and that VR estimation is particularly robust when viscosity contrast is close to background.

The quantitative error analysis in Fig. 3 reinforces this observation. Across the six evaluated material and geometry combinations, the lowest errors occurred for cases in which ER and VR were either moderate or clearly decoupled, while the highest ER errors occurred specifically for the two cases in which ER and VR were simultaneously large and equal, regardless of inclusion shape. This pattern indicates that prediction accuracy is not governed by the magnitude of ER or VR individually, but by the relationship between them, an effect not previously characterized for on-axis ARF-based viscoelastic estimation.

While these simulation results are encouraging, the simulated inclusions, though varied across three geometries, five focal depths, two F-numbers, and a broad range of material contrasts, were each mechanically homogeneous and isotropic within the inclusion itself. To demonstrate the framework beyond simulation, Fig. 4 shows that in an ex vivo chicken breast inclusion embedded in a gelatin background, the predicted ER and VR maps exhibit pronounced spatial heterogeneity with elevated values colocalized near one boundary of the inclusion rather than distributed uniformly as in simulated cases. This heterogeneity is consistent with the fibrous, anisotropic, internal structure of muscle tissue,inclusion property not represented in the training data, and suggests that, the model responds to genuine tissue structure rather than reproducing a fixed, training-like contrast pattern. Because ground-truth ER and VR were not independently measured, for this phantom, this result should be interpreted as evidence of plausible spatial generalization rather than validated accuracy.

In an in vivo murine tumor model, Fig. 5 and Fig. 6 showed that predicted ER and VR increased substantially from Day 2 to Day 33 in a control tumor (ER: 2.48 to 7.99; VR: 1.82 to 8.45), while a paclitaxel-treated tumor showed a comparably smaller increase over the same interval (ER: 5.51 to 7.04; VR: 3.05 to 5.68). Fig. 7 and Fig. 8 indicate that this pattern extended to the population level: control and paclitaxel groups exhibited similar median ER and VR at Day 4, but diverged progressively through Day 30–32, with control values consistently exceeding paclitaxel values at later timepoints. This divergence is consistent with a treatment-associated attenuation of the mechanical property changes that otherwise accompany tumor growth, supporting the relevance of the proposed ER and VR parameters to monitoring tumor progression and treatment response.

One limitation of the proposed framework is that it yields relative measures — elasticity ratio (ER) and viscosity ratio (VR) — rather than absolute moduli, as the normalization by background displacement features produces unitless contrast values that are meaningful only relative to an assumed homogeneous reference region within the same field of view. This makes the framework intrinsically better suited to focal, localized pathology such as tumors than to diffuse diseases where the entire tissue is uniformly altered and no such reference region exists. A related constraint is that the normalization step requires a mechanically homogeneous background to be accessible in the same acquisition; in tissue types where this assumption fails, the normalized features will not isolate mechanical contrast from excitation-induced spatial variation as intended.

Unlike QVisR [26], which employs domain adaptation through transfer learning on experimental phantom data to bridge the simulation-to-reality gap, the proposed pipeline was applied directly to experimental data without any fine-tuning or calibration on real tissue acquisitions. This may contribute to the tendency for predicted ER and VR values in the murine tumor to cluster at or beyond the upper bound of the training distribution’s highest contrast tier, and represents a concrete methodological gap relative to current best practice in the field.

Additionally, the network operates on seven hand-engineered displacement-derived features rather than the full displacement time-series used by QVisR [26] and shear-wave CNN methods. While this compact, interpretable input reduces computational cost and improves robustness to noise, any diagnostically relevant information in the temporal displacement profile not captured by these seven features is discarded before the network is applied.

Despite these limitations, this investigation demonstrates that the proposed framework is relevant to interrogating tissue in which localized elastic and viscous property changes are associated with pathology, such as tumor formation, without requiring per-A-line nonlinear model fitting or the depth- and elasticity-compensation procedures central to VisR. To implement the framework in practice, the following steps could be pursued: (1) ARFI data are acquired at a focal depth and F-number matching one of the configurations represented in the training dataset (Section 2.2); (2) displacement features are extracted and normalized to the same seven ratio features used in training; (3) the trained Stage 1 classifier is applied to generate an inclusion mask; (4) the trained Stage 2 regressor, using the predicted mask alongside the normalized features, produces parametric ER and VR images; and (5) these images are displayed for qualitative or, pending further validation, quantitative assessment of the region of interest.

This work has demonstrated the proposed pipeline for spherical, cuboidal, and ellipsoidal inclusions in simulation, an ex vivo chicken breast phantom, and an in vivo murine tumor model. Future work will pursue independent reference measurements of ER and VR for experimental validation, extend the training set to include anisotropic and heterogeneous inclusions representative of fibrous tissue structure, and evaluate the framework across a larger animal cohort to statistically confirm the treatment-related divergence observed here. Other potential applications of the framework, including renal and musculoskeletal assessment, are topics of ongoing investigation.

## 5 Conclusion

This work demonstrates that a two-stage machine learning pipeline can predict elasticity ratio (ER) and viscosity ratio (VR) at the A-line level directly from normalized ARFI-derived displacement features, without requiring per-A-line nonlinear MSD fitting or the depth- and elasticity-compensation procedures central to VisR. In silico, the pipeline accurately localized inclusion boundaries and recovered ER and VR contrast across spherical, cuboidal, and ellipsoidal geometries, with prediction accuracy shown to depend jointly on inclusion geometry and the relative magnitude of ER and VR, and with the most consistent performance observed when the two ratios were moderate or clearly decoupled. In an ex vivo chicken breast phantom, the trained model, without retraining, produced spatially bounded, contour-consistent ER and VR maps that captured internal tissue heterogeneity absent from the training data, indicating that the pipeline responds to genuine tissue structure rather than reproducing a fixed training-like pattern. In an in vivo murine tumor model, predicted ER and VR distinguished control from paclitaxel-treated tumors, with control tumors exhibiting a substantially larger increase in both parameters over the course of tumor growth than treated tumors, both at the representative-animal level and across the full imaged cohort. These results suggest that the proposed ER and VR parameters are relevant to assessing localized elastic and viscous property changes associated with tumor pathology and treatment response, and support the feasibility of a scalable, model-independent alternative to conventional VisR estimation for on-axis ARF-based viscoelastic imaging.

## Acknowledgment

This work was supported in part by a Cancer Center pilot grant funded through the NIH National Cancer Institute Cancer Center Support Grant P30CA071789, and by Dr. Hossain’s startup award from the University of Hawaii at Mānoa College of Engineering. The technical support and advanced computing resources from University of Hawaii Information Technology Services – Research Cyberinfrastructure, funded in part by the National Science Foundation CC* awards 2201428 and 2232862 are gratefully acknowledged.

## Notes

### Competing Interest Statement

The authors have declared no competing interest.

